# Natural pest control in African agriculture: can migratory birds offset the decline of native insectivores?

**DOI:** 10.64898/2026.11.17.767311

**Authors:** Ariane Dellavalle, Max D. Y. Khoo, Adam J. M. Devenish, Moses Chibesa, Jeff Mwila, Gabriel A. Jamie, Maggie Mwale, Frank J. Willems, Jan-Dieter Ludwigs, Joseph A. Tobias

## Abstract

Agricultural expansion and intensification are reducing populations of resident insectivorous birds worldwide, with potentially negative consequences for natural pest control, particularly in the tropics. Many migratory insectivores spend the non-breeding season in the tropics, often favouring disturbed habitats, potentially fulfilling ecological roles once played by resident species. However, few studies have examined the relative impacts of agricultural land-use change on resident and migratory bird species, particularly in the Afrotropics. Using field survey data from five study areas in Zambia, we examine how resident and migratory birds respond to land-use change and assess the contribution of long-distance migrants to invertivore functional diversity in human-modified landscapes. We show that, while resident invertivore richness and abundance decline, migrants are more species-rich and abundant in agricultural habitats than in forests. Trait-based analyses further reveal that migrants contribute unique functional diversity, particularly in smallholder and commercial farms, where their morphological traits overlap and extend beyond the current diversity of residents, occupying an estimated 44% of the total community morphospace in commercial farms. On one hand, our findings highlight the crucial importance of Afrotropical farmland for the conservation of migratory birds. On the other hand, the seasonal influx of migrants may have an important functional consequence, boosting the diversity and pest-predation potential of tropical bird communities. The conservation of migratory bird populations, throughout their breeding and wintering ranges, offers one route to supporting more sustainable agriculture in rapidly changing Afrotropical landscapes.

## 1. Introduction

Establishing sustainable and equitable food systems is a major goal for developing nations, particularly in sub-Saharan Africa where social inequality, human population growth and rising food demands present a host of social and environmental challenges (Leisher et al., 2022; Potapov et al., 2021). While socio-economic factors take centre stage, there is growing awareness that biodiversity could help to promote the transition to sustainable African food systems, for example through natural pest control, whereby predatory animals consume crop pests, boosting agricultural yields and reducing costs of pesticides for smallholder farmers (Garfinkel et al., 2022; Ocampo-Ariza et al., 2023; Olimpi et al., 2022). The potential for predators to provide this free service is being undermined through ongoing habitat loss and agricultural expansion driving rapid declines in native biodiversity, particularly in tropical regions (Cardinale et al., 2012; Song et al., 2018). However, few studies have quantified changes in predator populations in African agroecosystems, leaving many questions unanswered about the future of natural pest control and the types of land-use practices that could promote sustainable agriculture (Devenish et al., 2023; Seddon et al., 2016).

Birds are one of the main groups of predatory animals proposed to contribute to pest control (Bhoi et al., 2025; Mäntylä et al., 2011). Previous studies have shown that invertivorous birds (hereafter, ‘invertivores’) can play an important role in tropical agroecosystems by preying on crop pests, suppressing crop damage and boosting yields (Garfinkel et al., 2022; Jiménez-Albarral et al., 2026; Ocampo-Ariza et al., 2023; Olimpi et al., 2022). Unfortunately, resident invertivores are particularly sensitive to the removal of tropical forests and other natural habitats, undergoing widespread population declines in response to land-use change (Bregman et al., 2014; Sherry, 2021; Fig. 1a). Although this limits the insect predation services supplied by native invertivores in tropical agriculture (Bregman et al., 2016), a potential seasonal boost in supply of pest predation is provided by migratory invertivores that travel from high-latitude breeding ranges to the tropics during the non-breeding season. Pervasive population declines are also underway in many migratory invertivores for a wide array of reasons (Albert et al., 2020; Vickery et al., 2023), but a key difference with tropical invertivores is that migrants appear to prefer relatively disturbed tropical habitats during boreal winter (Levey et al., 2025). This raises the possibility that shortfalls in insect predation services in tropical agroecosystems can be at least partially offset by a periodic influx of migratory predators, if this coincides with crop growing seasons (Fig. 1a).

**Figure 1.**
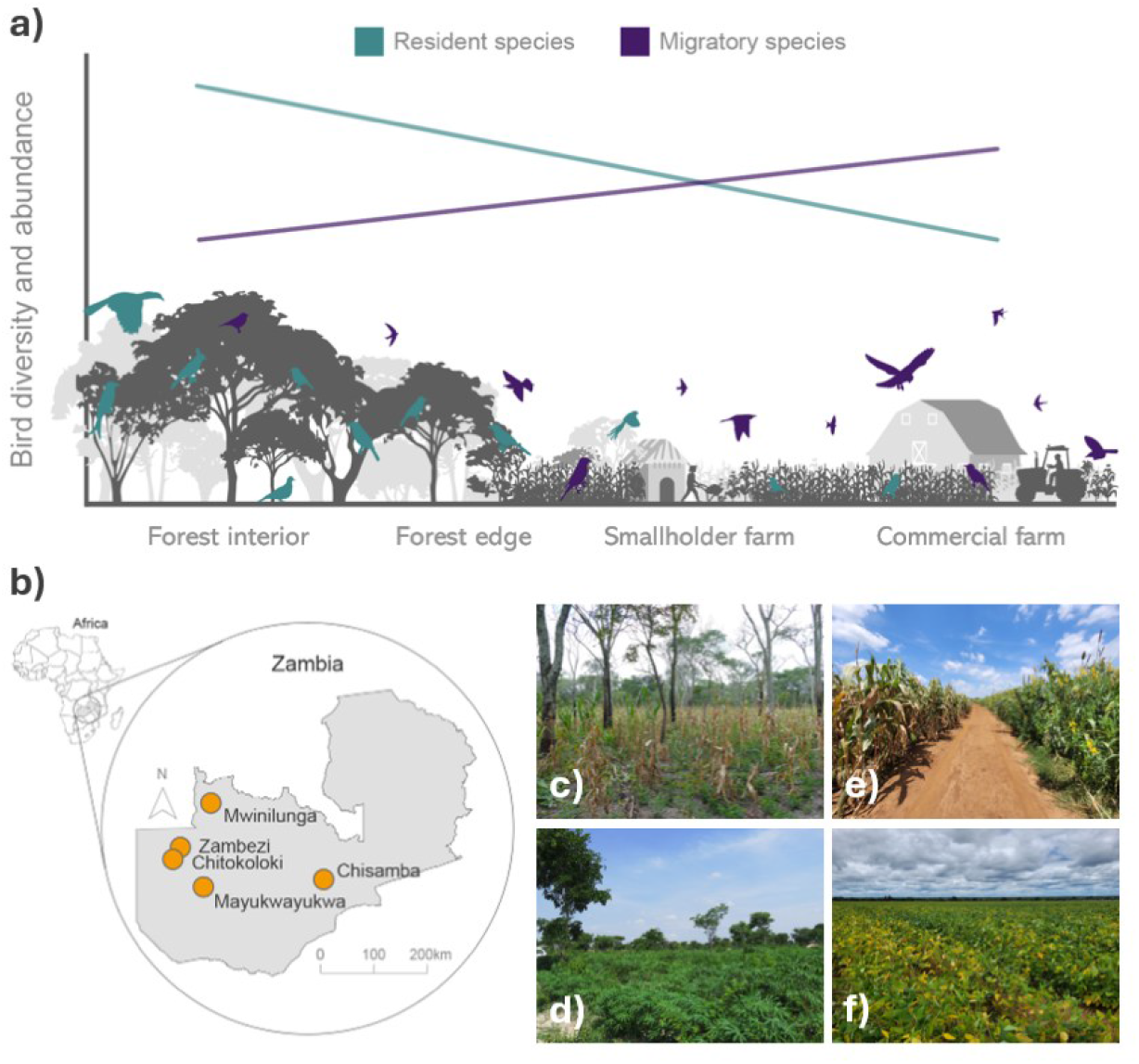
Potential impacts of land-use change on resident and migratory invertivorous birds and overview of the study system. (**a**) Conceptual framework illustrating the hypothesised effects of increasing land-use intensity from forest interior to commercial farming on bird abundance and diversity (including species richness and functional diversity). Abundance and diversity of resident invertivores is expected to decline, potentially reducing ecosystem functions and services such as arthropod pest control. In contrast, migratory invertivores may increase in diversity and abundance in human-modified landscapes owing to greater habitat suitability, altered resource availability or reduced competition, potentially offsetting these losses. **(b)** Location of the five study areas in Zambia. (**c-f**) Photographs showing examples of agricultural landscapes at study sites: smallholder cassava cultivation at Zambezi (**c**), smallholder maize cultivation at Mayukwayukwa with remnant native vegetation (**d**), and intensive commercial farming at Chisamba, including maize monoculture (**e**) and soybean production (**f**), with no native trees remaining in the landscape.

Increased awareness of the plight of long-distance migratory birds has focused conservation attention on protecting international flyways and habitat throughout the migratory cycle (Reynolds et al., 2017; Vickery et al., 2023). However, most of the attention has been on preserving suitable forest types without considering farmland or agroecosystem habitat, and our current understanding of the contribution of migratory birds to ecosystem services in agroecosystems is very sparse. Studies of avian ecosystem services rarely differentiate between migratory and resident species in temperate zone systems, while the focus of tropical studies is biased towards particular crops in South and Central America. Through a general review of literature, we identified 19 studies on ecosystem services provisioning by migratory birds in tropical agroecosystems, predominantly from the Neotropics (*n* = 13, 68.4%) and from coffee agroforests (*n* = 7, 36.8%), with only one study from the Afrotropics (Appendix A; Data S1). Therefore, the potential contribution of migratory birds to ecosystem services in tropical agriculture remains largely unknown, particularly in the Afrotropics (Greenler & Ebersole, 2015; Johnson et al., 2011; López-Hoffman et al., 2017).

To examine this question, we used point counts and mist-netting to survey resident and migratory invertivores across land-use gradients at five sites in Zambia (Fig. 1b). Zambia is a classic example of an Afrotropical country with rapid agricultural expansion and intensification driving widespread forest loss and land-use change (Denga et al., 2025; Ordway et al., 2017; Phiri et al., 2023). The agricultural landscape in Zambia, currently transitioning from forest and small-holder subsistence farming to large-scale commercial agriculture, offers a model system to explore the relationship between land-use change, biodiversity, and ecosystem function. In addition, many migratory bird species breeding in the Palearctic region (Europe, North Africa, and West, Central, and East Asia) migrate to Zambia during the wet season (November to April) which coincides with the growth and harvesting of most agricultural crops (Hachigonta et al., 2008).

By combining field survey data with functional trait data, we address two key questions related to bird populations in Zambian farmland. First, do migratory and resident invertivores differ in their use of anthropogenic compared to natural habitats? Second, to what extent does the seasonal presence of migratory invertivores make up for losses in species richness and functional diversity of resident invertivores? We assess the functional diversity of invertivore communities using comprehensive data on beak dimensions (Tobias et al., 2022), based on the assumption that beak morphology provides information on trophic niches, in particular the size and type of prey items selected (Bregman et al., 2016; Pigot et al., 2020; Tobias et al., 2020). In general, insectivore assemblages with a more diverse set of morphological traits and ecological roles will contribute to increased effectiveness in insect population control (Ceia et al., 2023; Philpott et al., 2009). Therefore, we interpret beak trait diversity as a proxy for the overall pest-predation potential of invertivore assemblages.

## 2. Methods

### 2.1 Study region and selection of study areas

We surveyed bird communities between 2020 and 2023 across five survey areas in central and north-western Zambia (Fig. 1b). The region has a subtropical climate, with a dry season from May to August and a wet season from September to April, annual rainfall of around 1,400 mm and a mean annual temperature of approximately 28 °C (Libanda et al., 2020).

North-western Zambia is characterised by dry evergreen forests dominated by *Cryptosepalum* trees, which are endemic to the region, and locally known as ‘Mavunda’, whereas central and northern Zambia are characterised by wet miombo woodlands (Chidumayo, 2019; Fanshawe, 2010). These forested areas are commonly cleared using a slash-and-burn technique for small-scale subsistence agriculture, primarily the cultivation of cassava and maize, on which approximately 75% of the local population relies (Food and Agriculture Organization of the United Nations, 2026). Under this shifting cultivation system, fields are often left fallow after a few years of cultivation, allowing for some degree of natural regeneration and creating a mosaic of forest, regrowth, and small crop fields (Masikati et al., 2021).

Within this landscape, we selected four study areas – Mwinilunga, Zambezi, Chitikoloki, Mayukwayukwa – to represent land-use gradients from mostly intact *Cryptosepalum* and miombo forests to secondary forests and smallholder agriculture (Fig. 1c-d). To capture the transition to larger-scale commercial farming, we selected a fifth study area, Chisamba, in central Zambia, where agricultural expansion and intensification have led to more extensive commercial farming operations, particularly of maize and soy (Fig. 1e-f).

### 2.2 Selection of survey sites and farms

In Mwinilunga, Zambezi, Chitikoloki, and Mayukwayukwa, we used a two-tier stratified random sampling design to select survey sites along a forest-to-shifting-agriculture gradient. First, each study area was overlaid with a spatially referenced 1 km × 1 km grid, dividing it into potential study sites. Grid cells were then classified into four broad land-use types based on forest-cover data from Hansen et al. (2013), accessed via Google Earth Engine (Gorelick et al., 2017). Given the mosaic landscape, we used an operational threshold of >95% forest cover to distinguish relatively intact and continuous forest (forest interior) from areas with greater land-cover modification, with remaining cells classified as forest edge (25-75% forest cover), smallholder agriculture (<5% forest cover) or other (e.g. urban areas or large water bodies). Only forest interior, forest edge and smallholder agriculture cells were retained for sampling. Second, 30% of grid cells in each eligible land-use category were randomly selected as survey sites for point counts and mist-netting. During fieldwork, site selection was adjusted where locations were inaccessible or community permissions could not be obtained. Survey transects followed existing paths or roads, with trails cut through the forest where necessary. Where vegetation, waterways or terrain prevented straight transects, curved or angled routes were used, minimising deviations where possible. In total, we selected 110 forest interior, 53 forest edge and 94 smallholder agriculture grid cells for surveying.

In Chisamba, we selected four commercial farms as sampling sites, each covering 2-10 km², and managed as a “conservation farm”. The main crops were soy and maize, while cover crops typically comprised mixtures of sunflower, millet, sunn hemp and sorghum. These farms used environmentally friendly practices including targeted pesticide application, cover cropping and crop rotation, which reduces reliance on fertilisers and other chemical inputs. This allowed us to investigate the potential of ecologically friendly commercial farming to support biodiversity and ecosystem services while reducing the confounding effects of agrochemical inputs when assessing the effects of land-use change.

### 2.3 Survey methods

In Mwinilunga, Zambezi, Chitokoloki and Mayukwayukwa, we surveyed birds between 2021 and 2022 over three field seasons, all conducted during the wet season (November-April). Surveys followed transects (Mwinilunga: *n* = 39; Zambezi: *n* = 30; Chitokoloki: *n* = 50; Mayukwayukwa: *n* = 52), each comprising 6-7 point-count sites at least 200 m apart to minimise double counting (Powell et al., 2015). In total, we surveyed 1,026 point-count sites across the four study areas (Mwinilunga: *n* = 229; Zambezi: *n* = 183; Chitokoloki: *n* = 302; Mayukwayukwa: *n* = 312). Counts lasted 10 minutes and were conducted between dawn and midday, when bird activity is typically highest. Observers remained quiet for one minute before recording all birds seen or heard, noting the species, abundance (recorded only during the second field season) and whether birds were using the vegetation within a 50-m-radius plot centred on the survey point. In total, these surveys produced 15,680 detections of 317 species. On commercial farms in Chisamba, we used survey methods broadly similar to those described above. Birds were surveyed between 2020 and 2022 over two field seasons (December-May and October-January, respectively). Depending on farm size, 2-10 survey points were selected per farm, spaced 1 km apart, for a total of 24 point-count sites. Each point was visited weekly, with a total of eight visits during the first season and nine during the second. Counts lasted 5 minutes, during which observers recorded all birds seen or heard and estimated the number of individuals in each cluster. In total, 634 surveys were conducted. After incorporating these data, our final dataset contained 19,294 detections of 377 bird species.

To assess whether the shorter survey duration on commercial farms (5 versus 10 minutes at other sites) affected our results, we conducted sensitivity analyses in which species and individual counts recorded on commercial farms were increased by 30%, 60% and 100% (Appendix B). These adjustments did not change our results, which is consistent with evidence that most individuals – typically over 90% – are detected within the first 5 minutes of a count, particularly in open habitats (Lee & Marsden, 2008; Mattos & Peris, 2008). We therefore retained the original counts and treated the surveys as comparable when grouping point-count sites into assemblage surveys.

To harmonise survey data across study areas, we averaged species richness (SR) and the number of resident and migratory invertivores (abundance) recorded at point-count sites across transects in Mwinilunga, Zambezi, Chitokoloki and Mayukwayukwa, creating a single assemblage survey within each 1 km^2^ grid cells. This reduced spatial autocorrelation between neighbouring points and enabled comparison with Chisamba, where a single point-count site was visited repeatedly within each 1 km² area. Averaging across point counts resulted in continuous values for SR and abundance. We retained continuous SR values to preserve variation in diversity, as rounding would reduce the number of distinct values. Abundance values were rounded to the nearest integer and retained as count data, as rounding had little effect given their higher values. For commercial farms in Chisamba, we obtained assemblage surveys by averaging data across repeated surveys at each site. Given the annual crop rotation practised on commercial farms, surveys conducted at the same site in different years were treated as independent sites. We restricted Chisamba data to November-April to align with surveys in the other study areas and to focus on the period when migrants overwinter, which coincides with the crop-growing season. Each resulting unit was referred to as an assemblage survey, and the bird community recorded within it as a species assemblage. In total, we obtained 217 assemblage surveys across four habitat types: forest (*n* = 77), forest edge (*n* = 32), smallholder farms (*n* = 60) and commercial farms (*n* = 48).

Point-count surveys can have low detection rates for smaller, less vocal and more cryptic bird species, particularly in dense vegetation (Mayhew et al., 2019; Wang & Finch, 2002). We therefore complemented point counts with mist-netting surveys to compare captures of resident and migratory invertivores across land-use types and to provide a complementary assessment of bird assemblage composition. We conducted mist-netting at 105 sites across the five study areas and four land-use types (forest: *n* = 32; forest edge: *n* = 42; smallholder farms: *n* = 22; commercial farms: *n* = 9). At each site, we deployed 80-300 m of mist nets, adjusting the total length to site configuration. Nets were 3 m high, with a mesh size of 30 mm. They were opened from dawn until midday to coincide with peak bird activity. Sampling was avoided during rain or strong winds. In total, we captured 1,949 individuals of 155 bird species, seven of which were not recorded during point-count surveys.

### 2.4 Species traits

Information on migratory behaviour, trophic niche and morphological traits was extracted from AVONET (Tobias et al., 2022) for all 383 species in our dataset. We classified each species as resident or migratory, defining migrants as species undertaking long-distance migrations and classifying species undertaking short-distance intratropical movements as residents. We restricted our analyses to invertivores, the main trophic guilds likely to feed regularly or opportunistically on invertebrate crop pests. To focus on potential pest predators, we excluded three invertivorous species considered unlikely to prey on crop pests: European Honey Buzzard (*Pernis apivorus*), Abdim’s Stork (*Ciconia abdimii*) and Hadada Ibis (*Bostrychia hagedash*).

Direct information on the diets of Zambian birds is extremely limited, so we used beak morphology as a proxy for avian trophic niches. Beak length, width and depth are associated with the size and type of food items consumed, as well as the foraging strategies used to obtain them (Pigot et al., 2016, 2020). We therefore selected these three traits to quantify functional diversity within invertivore assemblages. Variation in beak morphology may reflect differences in trophic niches and potential interactions with prey (Bregman et al., 2016; Pigot et al., 2020). We consequently interpreted greater beak trait diversity as an indicator of potentially broader pest-predation functions (Ceia et al., 2023; Philpott et al., 2009).

We measured beak length, width and depth data from all individual birds mistnetted and combined these with data published in AVONET, producing a data set of species means based on measurements of at least eight individuals per species (Tobias et al., 2022). To incorporate local morphological variation, we also measured these traits in birds captured during our mist-netting surveys in Zambia. Measurements from 1,933 individuals belonging to 155 species – including 1,153 individuals from 98 invertivore species – were added to AVONET, supplementing existing data with measurements from local populations.

### 2.5 Functional diversity of resident and migratory invertivores

Based on species-mean values for three beak measurements (length, depth and width), we quantified functional diversity as the volume of morphological trait space (hereafter, morphospace) using the function *hypervolume_gaussian* from the R package *hypervolume* v.3.1.6 (Blonder et al., 2014). For each assemblage (*n* = 217), we calculated the total morphospace volume occupied by all invertivores and the morphospace volume occupied by resident invertivores alone. Using these two diversity indices, we then calculated the proportion of morphospace occupied by resident invertivores relative to the total morphospace volume. Together, these metrics quantified the absolute morphospace volume of resident invertivores and the total functional diversity of each assemblage, providing a baseline against which to assess the contribution of migrants. We then quantified the additional morphospace contributed by migratory invertivores using the *hypervolume_set* function from the R package *hypervolume* (Blonder et al., 2014), which calculates the unique volume between two hypervolumes. This represents the volume of morphospace occupied by all invertivores but not by resident invertivores, thereby quantifying the additional trait space contributed by migrants (Appendix C). Finally, we calculated the proportion of total morphospace volume occupied by migrant invertivores. Together, these metrics allowed us to assess the extent to which migrants contribute to the functional diversity of invertivore assemblages beyond that represented by residents alone.

### 2.6 Statistical analyses

To assess whether migratory and resident invertivores differ in their use of anthropogenic versus natural habitats, we fitted generalised linear mixed-effects models (GLMMs), using land-use type (*n* = 4) – ranging from forest interior to commercial farms – as a categorical predictor and SR and abundance as response variables. Models were fitted separately for residents and migrants. As SR was averaged across point counts within each assemblage survey (*n* = 217), resulting in continuous values, SR was modelled using a Gaussian distribution. Abundance values were rounded to the nearest integer and modelled using a generalised Poisson distribution to account for overdispersion.

To assess how the contribution of migratory invertivores to trophic functional diversity – as a proxy for pest-predation potential – varied with land use, we fitted four GLMMs using morphospace volume metrics as response variables. These comprised the morphospace volume occupied by residents, the additional morphospace contributed by migrants, the proportion of total morphospace occupied by residents, and the proportion represented by the additional morphospace contributed by migrants. Resident morphospace volume and additional migrant morphospace volume were log-transformed and modelled using Gaussian distributions. The proportions of morphospace occupied by residents and contributed by migrants were modelled using beta distributions.

All models were fitted using the *glmmTMB* v. 1.1.11 R package (Brooks et al., 2017), with field season included as a predictor to account for seasonal variation in bird assemblages and migration patterns. We assessed model fit using simulated residual diagnostics and examined whether residual patterns differed across land-use types and seasons. Where diagnostics indicated variation in residual dispersion, we fitted alternative models allowing dispersion to vary with season, land-use type, or both, and compared model fit using Akaike’s information criterion (AIC). The final resident morphospace volume model allowed dispersion to vary with season, whereas the models of the proportions of morphospace occupied by residents and contributed by migrants allowed dispersion to vary with both land-use type and season. The additional migrant morphospace volume model retained constant dispersion. We performed post hoc pairwise comparisons between land-use types using the *emmeans* v.1.11.1 R package, applying Holm adjustments for multiple comparisons (Lenth et al., 2024; Searle et al., 1980). For models of morphospace proportions, estimated marginal means were back-transformed from the logit scale to the response scale using *emmeans* to obtain percentage contributions. We carried out all statistical analyses in R v.4.5 (R Core Team, 2018).

## 3. Results

Surveys yielded 19,294 point-count observations and 1,949 mist-net captures, for 383 bird species, of which 198 were invertivores. Total pooled species richness (SR) was broadly similar across land uses, with the highest richness recorded at forest edges. This pattern reflected higher richness of granivores, omnivores and vertivores at forest edges (Fig. 2a). A similar pattern was observed on commercial farms, although richness of invertivores was lower (Fig. 2a). Of the 383 species, 340 were residents and 43 migrants; all migrants were predators, 31 (72%) of which were invertivores (Fig. 2b).

**Figure 2.**
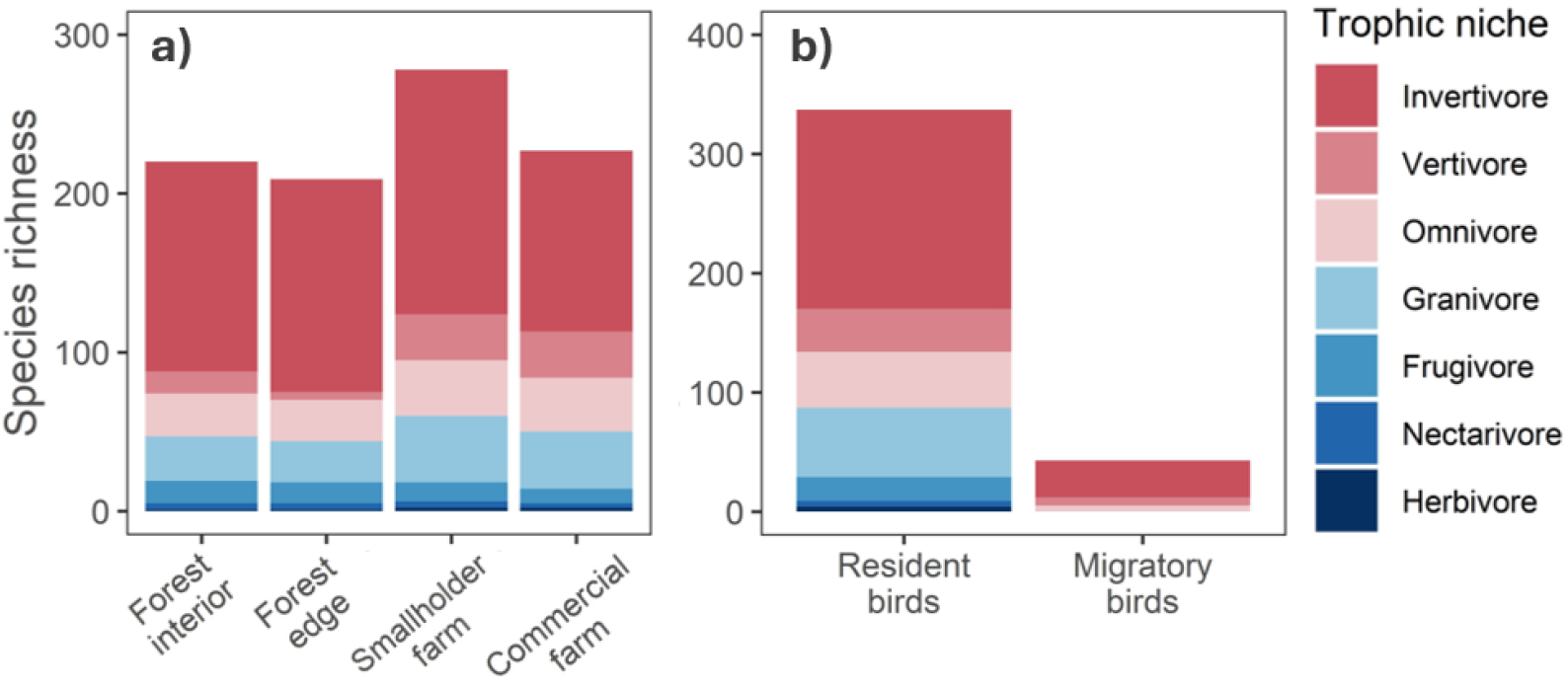
Trophic structure of avian assemblages partitioned by land use and migratory behaviour. Bar plots show the number of bird species within trophic niche across (**a**) land-use types and (**b**) migratory categories. Species richness estimates are based on point-count and mist-net surveys. Colours indicate primary trophic niche, extracted from a global database (Tobias et al., 2022), highlighting the prevalence of invertivory among migrants.

### 3.1 Effects of land use on resident and migratory invertivores

Resident invertivore SR and abundance did not differ significantly between forest interior and forest edge (Fig. 3a,c; Appendix D). However, resident SR was significantly lower in smallholder farms than in forest interior (17% decrease; estimated means = 4.02 and 3.34, respectively; *p* < 0.0001) and forest edge (15% decrease; estimated means = 3.93 and 3.33, respectively; *p* < 0.01), whereas abundance did not differ significantly between smallholder farms and either forest habitat. Both SR and abundance were significantly lower in commercial farms than in all other land-use types (Fig. 3a,c; Table 1; Appendix D). The largest decline in SR was between forest interior and commercial farms, where estimated mean SR was 80% lower (4.02 vs 0.79 species per assemblage survey; *p* < 0.0001).

**Figure 3.**
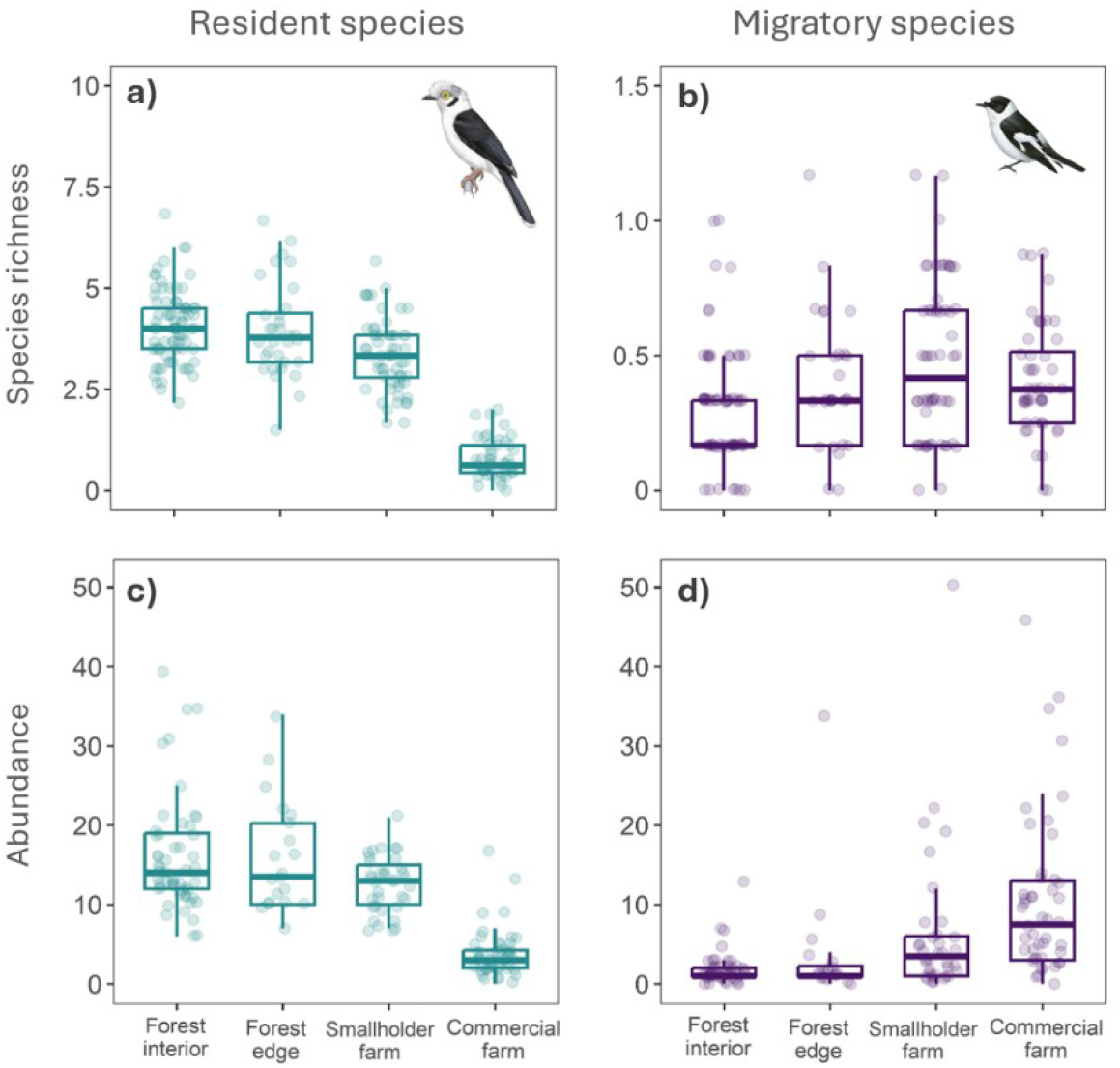
Contrasting responses of resident and migratory invertivores to land-use intensification. Boxplots show species richness (**a-b**) and abundance (**c-d**) of resident (**a,c**) and migratory (**b,d**) invertivorous birds across four land uses. Species richness and abundance were calculated by pooling observations across spatial and temporal replicates and dividing by the total number of replicates, yielding the mean number of species and individuals per survey point. Jittered points show these mean values for individual survey points. Illustrations show example species: *Prionops plumatus* (tropical resident) and *Ficedula albicollis* (long-distance migrant).

**Table 1.** Impact of land use on the species richness and abundance of resident and migratory invertivores. Results shown are model summaries predicting the effect of land use on bird response groups. Statistically significant effects are indicated in bold. Effect sizes have not been back-transformed. Details on model structure are described in Methods; Holm-adjusted pairwise comparisons among land uses are reported in Appendix D.

| **Coefficient** | **Species richness** |  |  | **Abundance** |  |  |
|---|---|---|---|---|---|---|
|  | **Estimate ± SE** | **z-value** | **p-value** | **Estimate ± SE** | **z-value** | **p-value** |
| **Resident invertivores** |  |  |  |  |  |  |
| Intercept (Forest) | 4.024 ± 0.14 | 29.58 | < 0.0001 | 2.759 ± 0.05 | 54.84 | < 0.0001 |
| Forest edge | -0.086 ± 0.18 | -0.48 | 0.631 | 0.001 ± 0.09 | 0.01 | 0.993 |
| Smallholder farm | **-0.683 ± 0.15** | **-4.65** | **< 0.0001** | **-0.197 ± 0.08** | **-2.36** | **< 0.05** |
| Commercial farm | **-3.231 ± 0.21** | **-15.68** | **< 0.0001** | **-1.393 ± 0.12** | **-12.01** | **< 0.0001** |
| **Migratory invertivores** |  |  |  |  |  |  |
| Intercept (Forest) | 0.268 ± 0.03 | 9.45 | < 0.0001 | 0.983 ± 0.23 | 4.16 | < 0.0001 |
| Forest edge | 0.098 ± 0.05 | 1.88 | 0.061 | 0.214 ± 0.27 | 0.78 | 0.434 |
| Smallholder farm | **0.199 ± 0.04** | **4.65** | **< 0.0001** | **0.767 ± 0.21** | **3.67** | **< 0.001** |
| Commercial farm | **0.151 ± 0.05** | **3.26** | **< 0.01** | **1.326 ± 0.27** | **4.85** | **< 0.0001** |

In contrast, migratory invertivore SR and abundance were significantly higher in smallholder and commercial farms than in forest interior. SR increased by 86% in smallholder farms (*z*-ratio = 4.65; *p* < 0.0001) and 65% in commercial farms (*z*-ratio = 3.26; *p* < 0.01) relative to forest interior. Conversely, abundance increased most in commercial farms, rising by 135% (*z*-ratio = 4.85; *p* < 0.0001) compared with 78% (*z*-ratio = 3.67; *p* < 0.01) in smallholder farms relative to forest interior. Although migrant SR did not differ significantly between forest edge and either farm type, migrant abundance was significantly higher in commercial farms than at forest edges (Fig. 3b,d; Table 1; Appendix D).

Mist-net captures reflected the pattern observed in assemblage surveys, with the representation of migrants increasing with land-use intensification. Migrants accounted for just 1% and 2% of captures in forest interior and forest edge, respectively, with three and four migratory species recorded (Fig. 4a-b). Their representation increased markedly in agricultural habitats, comprising 15% of captures in smallholder farms and 47% in commercial farms, where eight migratory species were captured (Fig. 4c-d).

**Figure 4.**
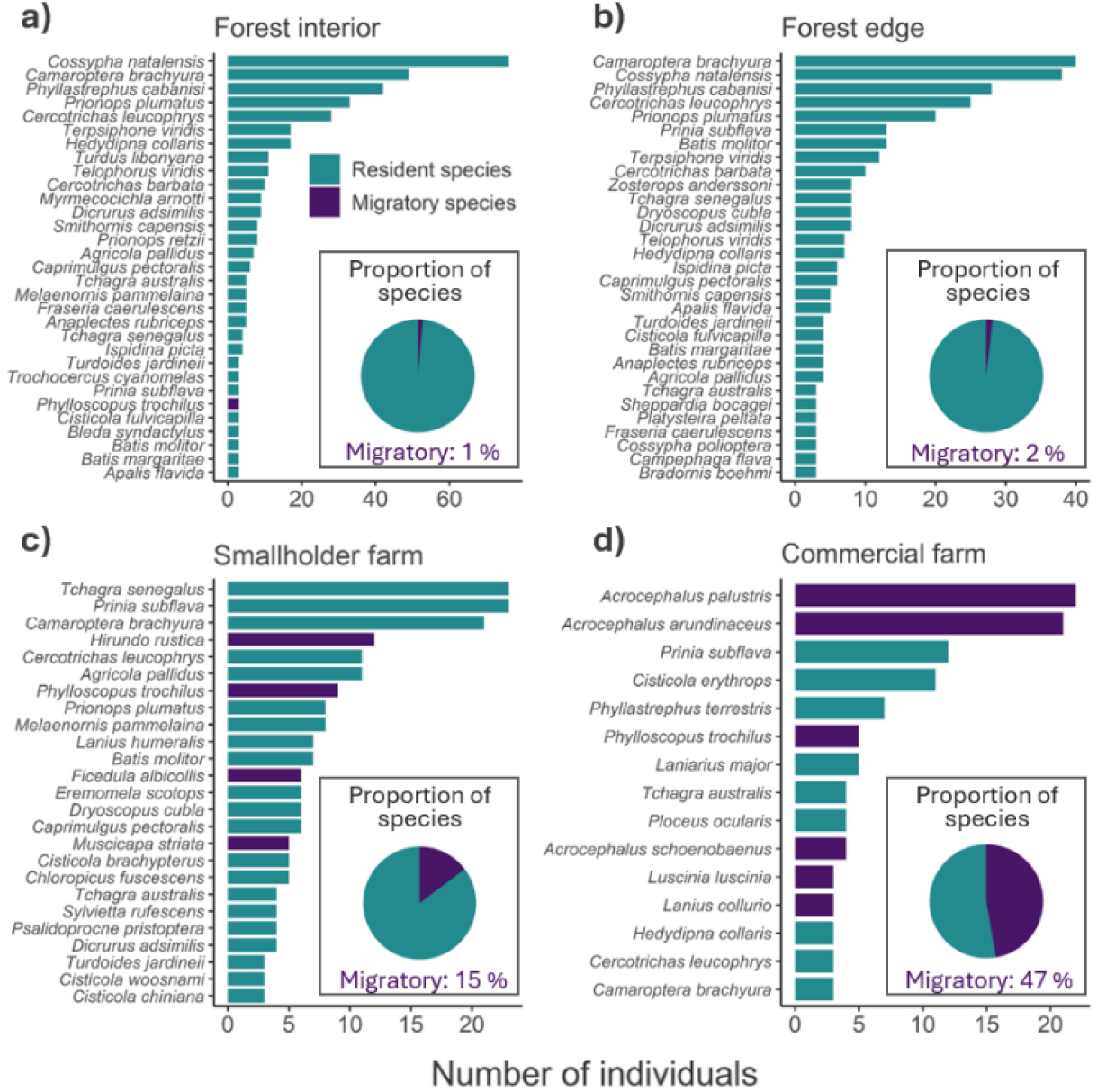
Relative abundance of resident and migratory invertivores captured in mist nets across different land-uses. Horizontal bars show the total number of individuals captured for each invertivorous bird species in: **(a)** forest interior, **(b)** forest edge, **(c)** smallholder farms, and **(d)** commercial farms. Pie charts indicate the proportion of resident and migratory species recorded in each land use. Species represented by fewer than two individuals within a habitat were omitted from the bar charts for visualisation purposes.

### 3.2 Migratory invertivores increase functional diversity in anthropogenic habitats

Given the contrasting patterns in resident and migratory invertivore SR and abundance across land uses, we next assessed whether beak-trait diversity, as a proxy for pest-predation potential, showed similar patterns and the extent to which migrants contributed additional functional diversity. Absolute morphospace volume occupied by resident invertivores did not differ significantly among forest interior, forest edge and smallholder farms, but was estimated to be 67.5% lower on commercial farms than in forest interior, although this difference was marginally non-significant after Holm correction (uncorrected *p* = 0.012; adjusted *p* = 0.052; Fig. 5c; Table 2; Appendix D). The proportion of total morphospace occupied by resident invertivores was also significantly lower on commercial farms than in forest interior (*z*-ratio = 4.77; *p* < 0.0001), forest edge (*z* = 3.67; *p* < 0.01) and smallholder farms (*z* = 4.32; *p* < 0.001), with no significant differences among these three land-use types (all adjusted *p* > 0.8; Fig. 5e; Table 2; Appendix D).

**Figure 5.**
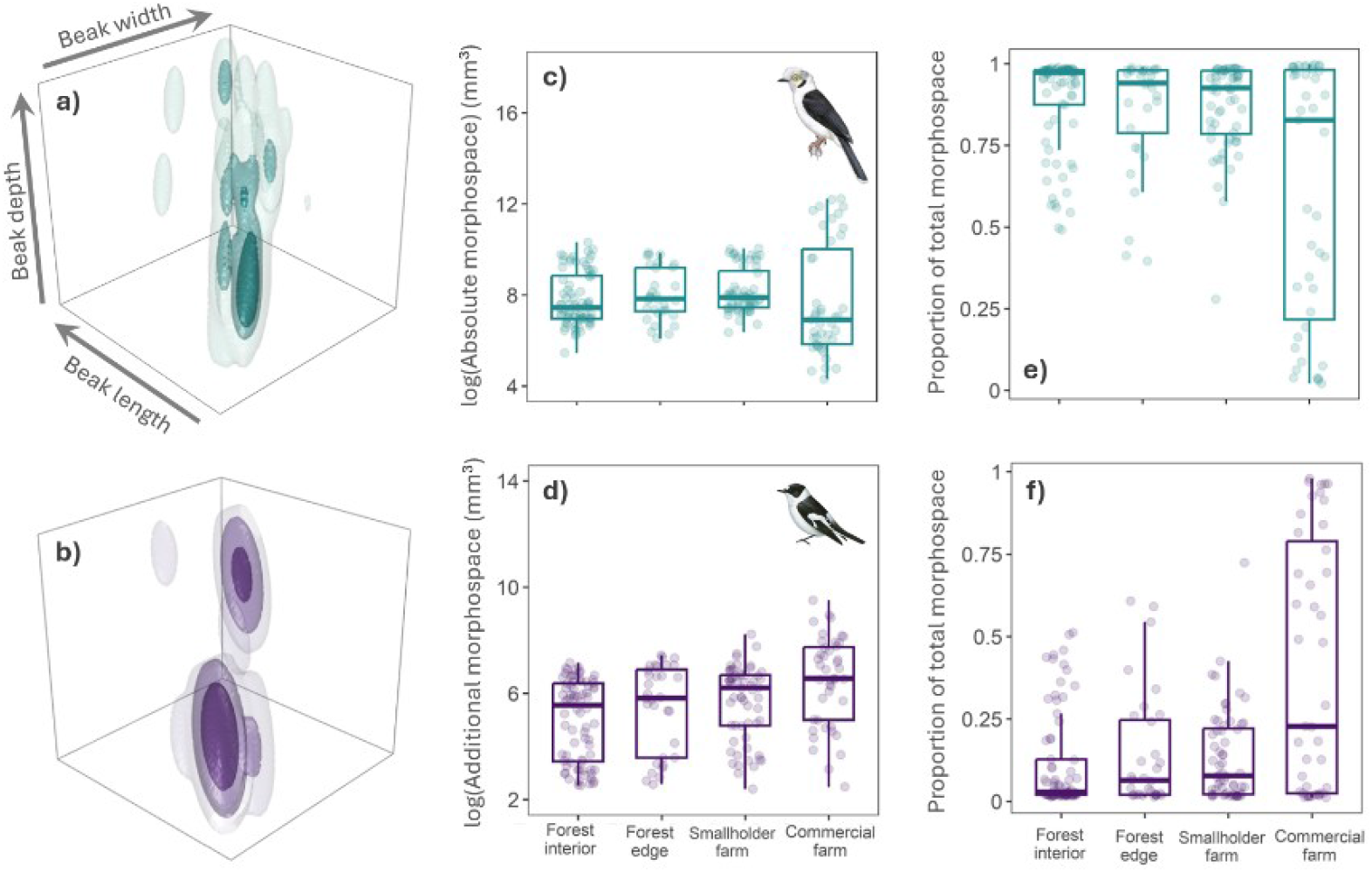
Contribution of resident and migratory invertivores to the functional diversity of bird assemblages across a land-use intensity gradient. Functional trait distributions are shown for (**a**) resident and (**b**) migratory species. Colours indicate species density in a multivariate trait space (‘morphospace’) defined by three beak traits associated with feeding: length, width and depth. Isopleths enclose 90%, 50% and 25% of the estimated species distribution in each land use. Arrows in (a) indicate increasing values of each beak trait. The contributions of residents and migrants to morphospace volume are shown as (**c**) the absolute volume occupied by residents, (**d**) the additional volume contributed by migrants beyond that occupied by residents (Appendix C), and the proportion of total volume occupied by (**e**) residents and (**f**) migrants. Jittered points represent mean morphospace volume metrics per survey point, calculated across spatial and temporal replicates. Illustrations representing resident and migratory species depict *Prionops plumatus* (resident) and *Ficedula albicollis* (migratory), respectively.

**Table 2.** Impact of land use on the morphospace contribution of resident and migratory invertivores. Results shown are model summaries predicting the effect of land use on four morphospace metrics: absolute volume occupied by residents, additional volume contributed by migrants, and the proportions of total volume occupied by each group (Figure 5; Appendix C). Statistically significant effects are indicated in bold. Effect sizes have not been back-transformed. Details on model structure are described in Methods, and Holm-adjusted pairwise comparisons among land uses are reported in Appendix D.

| Response variable | Response group | Land use | Estimate ± SE | z-value | p-value |
|---|---|---|---|---|---|
| **Absolute morphospace** | **Resident invertivores** | Intercept (Forest) | 7.36 ± 0.28 | 26.34 | < 0.0001 |
|  |  | Forest edge | 0.108 ± 0.23 | 0.48 | 0.632 |
|  |  | Smallholder farm | 0.27 ± 0.19 | 1.44 | 0.149 |
|  |  | Commercial farm | **-1.123 ± 0.45** | **-2.51** | **< 0.05** |
| **Additional morphospace** | **Migrant invertivores** | Intercept (Forest) | 4.624 ± 0.41 | 11.22 | < 0.0001 |
|  |  | Forest edge | 0.519 ± 0.33 | 1.6 | 0.111 |
|  |  | Smallholder farm | **0.741 ± 0.27** | **2.78** | **< 0.01** |
|  |  | Commercial farm | **1.162 ± 0.5** | **2.31** | **< 0.05** |
| **Morphospace proportion** | **Resident invertivores** | Intercept (Forest) | 1.832 ± 0.31 | 5.87 | < 0.0001 |
|  |  | Forest edge | -0.251 ± 0.25 | -1.01 | 0.315 |
|  |  | Smallholder farm | -0.124 ± 0.19 | -0.64 | 0.52 |
|  |  | Commercial farm | **-1.755 ± 0.37** | **-4.69** | **< 0.0001** |
|  | **Migrant invertivores** | Intercept (Forest) | -1.827 ± 0.31 | -5.9 | < 0.0001 |
|  |  | Forest edge | 0.259 ± 0.25 | 1.05 | 0.295 |
|  |  | Smallholder farm | 0.166 ± 0.19 | 0.87 | 0.385 |
|  |  | Commercial farm | **1.819 ± 0.39** | **4.93** | **< 0.0001** |

In contrast to the patterns observed for resident invertivores, the additional morphospace contributed by migrants was significantly greater on smallholder farms than in forest interior (*z*-ratio = 2.78; *p* < 0.05), with estimated unique morphospace volume approximately 110% higher. Although estimated additional morphospace volume was also higher on commercial farms (approximately 220% higher than in forest interior), this difference was not significant after Holm correction (uncorrected *p* = 0.021; adjusted *p* = 0.109; Fig. 5d; Table 2; Appendix D). No significant differences were found between forest edge and forest interior or among the other land-use types (all adjusted *p* > 0.7; Appendix D). However, migrants occupied a significantly greater proportion of total morphospace on commercial farms than in forest interior (*z*-ratio = 5.03; *p* < 0.0001), forest edge (*z*-ratio = 3.88; *p* < 0.001) and smallholder farms (*z*-ratio = 4.48; *p* < 0.0001; Fig. 5f; Appendix D).

## 4. Discussion

Our surveys detected contrasting responses of resident and migratory invertivores to increasing land-use intensity across a gradient from natural forests and smallholder farming to large-scale commercial agriculture. While resident species richness and abundance declined, migratory invertivores became more abundant and species-rich in agricultural habitats. We also show that this influx of migrants contributes unique functional space to agricultural bird assemblages, while also occupying the functional space occupied residents. This combination of functional complementarity and overlap suggests that migratory birds may both extend the range of trophic niches represented in agricultural habitats and reinforce existing ecological roles by providing an ecological back-up to declining native species. These findings highlight the importance of Afrotropical agriculture as key habitats maintaining populations of long-distance migrants that are generally declining in Europe and Asia (Vickery et al., 2023). In addition, they suggest that seasonal migrant populations may play a pivotal role in maintaining functional diversity of potential pest-predators in and adjacent to cropland and agroforestry, with implications for the resilience and sustainability of agricultural production landscapes in Africa.

### 4.1 Contrasting land-use change responses of resident and migratory invertivores

The contrasting responses of resident and migratory invertivores to land-use change may reflect differences in their movement patterns, particularly in their ability to cross gaps in habitat and cope with seasonal variation in resource availability (Weeks et al., 2022). While migratory species can move across landscapes and between regions to exploit seasonal resource peaks, resident invertivores are highly sedentary, usually remaining in the same home range or territory throughout the year (Salisbury et al., 2012; Tobias et al., 2013). Their relatively sedentary behaviour and reduced morphological adaptation for dispersal make resident tropical insectivores particularly vulnerable to the loss, degradation and fragmentation of natural vegetation (Weeks et al., 2023).

The declines in tropical forest invertivores following habitat modification and agricultural intensification are well documented (e.g. Bregman et al., 2016; Jarrett et al., 2021; Sekercioglu et al., 2002). In our study, declines in resident invertivores were particularly pronounced on commercial farms, whereas smallholder farms retained relatively high resident diversity, with abundance not differing significantly from forest habitats. This may reflect the greater structural heterogeneity of smallholder landscapes, where a mosaic of cultivated fields, fallows, trees and remnant natural vegetation provides a wider range of foraging and nesting opportunities. The cultivation of different crops, including annual and perennial species such as maize and cassava, may also provide a more varied supply of resources throughout the year (Johnson et al., 2011). Our findings therefore support previous studies showing that heterogeneous smallholder farming systems can retain substantial diversity of forest-associated birds within agricultural landscapes (e.g. Jarrett et al., 2021; Rurangwa et al., 2022; Šálek et al., 2026).

Palearctic migrants occur in Zambia during their non-breeding period which coincides with the wet-season, when peak rainfall supports both crop growth and increased invertebrate activity. The resulting availability of insects in cultivated fields and associated vegetation could provide important foraging opportunities for migrants, particularly where prey is more readily accessible than in forest habitats (Vickery et al., 2023) or competition with resident invertivores is lower (Cox, 1985). The high mobility and resource-tracking behaviour of certain migratory invertivore species also allow them to actively move within their wintering grounds to maximise their access to food sources (Koleček et al., 2018; Thorup et al., 2017). In addition, many migratory species recorded in our study are associated with open habitats (Vickery et al., 2023), and the vegetation structure of crops such as maize may resemble that of natural open or wetland habitats used by these species (Fontanilles et al., 2020; Galle et al., 2009), making agricultural landscapes suitable for foraging. The presence of great reed warblers (*Acrocephalus arundinaceus*) and marsh warblers (*Acrocephalus palustris*), which were among the most frequently recorded species in low-stature cover crops (Fig. 4d), is consistent with this possibility.

It is important to note that the high diversity of migrants recorded in agricultural habitats may partly reflect the management practices of the farms included in our study. In particular, the commercial farms surveyed were relatively low-input systems, incorporating practices such as cover cropping and reduced pesticide use. These practices may help maintain invertebrate resources and vegetation structure, potentially contributing to the suitability of these farms for insectivorous birds (Dänhardt et al., 2010; Hole et al., 2005). Whether similar patterns would occur on conventional, high-input commercial farms remains uncertain.

### 4.2 Migratory birds expand functional diversity of potential pest-predators in agricultural lands

The functional trait space occupied by migratory insectivores on Zambian farmland overlapped broadly with the functional roles of resident species. Migrants may therefore provide functional redundancy in trophic niches, increasing the number of species and individuals supplying a pest-control service, and thereby boosting the functional resilience of ecosystem functions to further shocks such as land-use intensification, climate change and disease (Weeks et al., 2025). As well as filling gaps within resident functional space, we also found that migratory species increased the overall functional diversity of agricultural bird assemblages by adding beak morphologies not represented among resident species (a proxy for the number of insect-predating niches occupied by birds). This expansion of functional trait space was greatest in smallholder farms, while the proportional increase in overall trait space provided by resident species was greatest in commercial farms.

These findings have potential implications for the pest-predation services provided by invertivorous birds. A greater diversity of beak morphologies may allow bird assemblages to exploit a wider range of prey sizes and types (Philpott et al., 2009; Pigot et al., 2020; Schleuning et al., 2023), potentially increasing the range of crop pests subject to predation. Migrants may be particularly important in this respect because their presence coincides with the growing season in Zambia, when crops and associated invertebrate populations are active. Previous work in Neotropical coffee agroforests has linked greater migratory bird diversity with lower invertebrate densities, providing some support for the possibility that migrants contribute to pest regulation in tropical agriculture (Bael et al., 2008). In our study, however, we did not directly measure prey consumption, pest abundance or crop damage. The contribution of migrants to pest control seems very likely given the expansion in population and functional diversity of insectivores on Zambian farmland during the crop growing season. However, this hypothesis requires further investigation with direct observations, experimentation and dietary analysis.

The potential contribution of migrants to agroecosystem services also has important conservation implications. Many migratory invertivores using Zambian farmland breed in the Palearctic and depend on suitable habitats across multiple countries and along their migratory routes. Protecting these species therefore requires coordinated conservation across their annual cycle, including breeding habitats, migratory stopover sites and non-breeding habitats in the Afrotropics (Vickery et al., 2023). Maximising the suitability of Afrotropical farmland as bird habitat may therefore have a double benefit of promoting conservation of large migratory bird populations, while also increasing the potential pest-predation services provided by avian insectivores in Europe as well as Africa (Altieri, 2004; Larsen & Noack, 2020).

The potential benefits of migratory birds to farmers in Zambia highlights how conservation action in developed nations at higher latitudes can have positive implications for agricultural systems in the tropics. However, the value of agricultural habitats to migrants will depend on how they are managed. The relatively high representation of migrants on the commercial farms surveyed here may partly reflect their comparatively low pesticide use and the presence of cover crops. Our study calls for a suite of agricultural management measures to be implemented in Africa and beyond. Commercial farms should be encouraged to establish a mixture of crops and crop-rotation to provide more heterogenous landscapes, boosting bird populations (Priyadarshana et al., 2021). The success of Chisamba eco-farms suggests that regular cover cropping and reduced pesticide load will provide rich habitat for long-distance migrant birds while also increasing the potential of the local bird assemblage to provide nature-based solutions to crop damage by insects and other pests. These interventions should be focused towards regions of high importance for migratory insectivores and integrated into wider trade-offs between maintaining biodiversity and agricultural productivity in the Afrotropics (Chapman et al., 2026).

### 4.3 Limitations and future directions

Our survey methods may have underestimated migrant diversity. Point counts are less effective at detecting some small, cryptic or less vocal species (Robinson et al., 2018; Wang & Finch, 2002), and our mist-netting data from commercial farms revealed several migratory insectivores that were either not recorded or were less frequently detected during point counts. These included species such as Thrush Nightingales (*Luscinia luscinia*), Common Whitethroats (*Curruca communis*) and Reed Warblers, which forage within vegetation and may consume invertebrates directly from crops. Such species could be particularly relevant to the consumption of crop pests, including Fall Armyworm (*Spodoptera frugiperda*), which has caused substantial agricultural damage in Zambia (Day et al., 2017; Kansiime et al., 2019; Tambo et al., 2020). More comprehensive surveys, combined with dietary analyses, will be important for establishing which migratory species consume crop pests and the extent to which they contribute to pest regulation.

Our sampling methods are relatively simplistic, lacking explicit detection modelling and pooling datasets with slightly different methods. For example, the inclusion of point counts with different durations may introduce some biases in estimates of species richness and abundance. However, we used a sensitivity analyses (Appendix B) and parallel surveys with mistnets (Fig. 4) to show that this does not alter our main conclusions. In addition, our use of beak morphology as a proxy for function indicates potential prey use but does not directly measure diet or pest control.

Future studies should focus on exploring three main research avenues. First, the low-input commercial farms we surveyed do not represent conventional commercial agriculture, so targeted comparisons are needed to isolate the effects of individual agricultural practices (e.g. low vs high pesticide use; use of diverse cover crops). This type of detailed comparison across farming systems and management practices are needed to develop more specific guidelines for nature-based solutions in Afrotropical agriculture. Second, we did not quantify land-use history, proximity to natural forest and habitat connectivity, all of which may influence bird communities. Further research accounting for these factors is required to help distinguish the effects of local agricultural management from those of the surrounding landscape. Finally, direct tests are needed to help establish the importance of pest-predation services by resident and migratory birds, and the effects on pest populations and crop yields. Methods such as faecal DNA metabarcoding, bird-exclusion experiments, and estimates of leaf-damage and crop yield, would help to clarify the role of birds and other wildlife in supplying cost-effective pest-predation services (Priyadarshana et al., 2021).

### 4.4 Conclusion

Given the global rise in food demand, and projected increases in pest damage due to climate change (Deutsch et al., 2018; Hatfield et al., 2020; Leisher et al., 2022), understanding the pest-regulating potential of animal diversity within agroecosystems is an important first step towards promoting appropriate nature-based solutions. Our surveys reveal that resident invertivore species richness, abundance and functional diversity is lower in commercial farms compared with other habitats, with an opposite trend found in migratory invertivores which are present for part of the year. Although there is a net decline in species richness, abundance, and functional diversity of invertivores in commercial farms compared to forests, this seasonal influx of long-distance migrants at least partially offsets the decline. As shown in commercial shade coffee plantations in Central and South America (e.g. Greenberg et al., 1997), Afrotropical agriculture may provide an important refuge for migratory bird species. In this case, small-holder agriculture or low-input commercial cropland may offer highly suitable habitat for a subset of species, perhaps as a direct result of reduced competition with resident invertivores which have been largely removed from more intensive agricultural landscapes by deforestation. Migratory species could therefore play a major role in boosting natural pest control services that assist farmers by reducing the costs associated with pesticide application, potentially making agriculture more sustainable. To maintain high-density populations of migratory bird species, agricultural land-use strategies in sub-Saharan Africa should focus on maintaining crop heterogeneity, promoting adjacent patches of natural habitat, reducing pesticide use, and planting mixed cover-crops on fallow fields. Equally, conservation efforts focused on maintaining breeding populations of long-distance migratory birds on their breeding grounds in Europe and Asia could ultimately help to support sustainable farming in the Afrotropics. Further work is needed to disentangle the role of migratory and resident invertivores as nature-based solutions to crop pests, and to understand the functional role of migratory birds in agroecosystems more generally.

## Author contributions

**AD:** Data curation, Formal analysis, Investigation, Visualisation, Writing – original draft, Writing – review and editing. **MDYK:** Conceptualization, Data curation, Formal analysis, Investigation, Methodology, Visualisation, Writing – original draft, Writing – review and editing. **AJMD:** Conceptualization, Investigation, Methodology, Project administration, Supervision, Writing – review and editing. **MC:** Investigation, Methodology, Project administration. **JM:** Investigation, Methodology. **GAJ:** Investigation, Methodology. **MM:** Investigation, Methodology. **FJW:** Investigation, Methodology. **JDL:** Conceptualization, Funding acquisition, Investigation, Methodology, Project administration. **JAT:** Conceptualization, Funding acquisition, Investigation, Methodology, Project administration, Supervision, Visualization, Writing – original draft, Writing – review and editing.

## Data accessibility statement

The data that support the findings of this study are openly available in Helix at https://helix.imperial.ac.uk/xxxx (full link to be added on acceptance)

## Declaration of AI use

The free version of OpenAI’s ChatGPT (GPT-5 and GPT-6) was used as a search engine, to provide suggestions for debugging code, to proofread individual paragraphs, and to suggest alternative ways of phrasing sentences.

## Acknowledgements

We thank Mike Ford, David Chapoloko, Dui Jasinghe, Alex Rabeau, and Leslie Reynolds for help with fieldwork, data or useful discussions. This research was funded by the UKRI Global Challenges Research Fund grant ES/P011306/1 (JAT). AD was supported by a Natural Environment Research Council studentship through the Science and Solutions for a Changing Planet Doctoral Training Partnership. MDYK was supported by the Singapore Ministry of National Development EDGE scholarship.

## SUPPLEMENTARY MATERIALS

As separate Excel file: **Data S1**

**Sheet 1:** Metadata. Contains descriptions and definitions for all variables.

**Sheet 2:** Survey data. Contains point-count data including background information on survey-point location, season and land-use type, together with the full list of species and relative abundance at each survey point during field surveys.

**Sheet 3:** Mist-net data. Contains mist-net capture data including background information on mist-net site location, season and land-use type, together with the full list of species and number of captures recorded at each mist-net survey site.

**Sheet 4:** Traits. Contains a matrix of species traits, including migratory behaviour, trophic niche and beak morphological traits.

**Sheet 5:** Literature survey. Contains a full list of studies identified in the literature that examine the role of migratory birds in providing ecosystem services.

## Appendix A. Systematic review of existing studies on ecosystem services provided by migratory birds in agroecosystems

We conducted a systematic review to identify studies investigating ecosystem services provided by migratory birds in agroecosystems, focusing on their wintering ranges. On 19 October 2026, we searched Web of Science for peer-reviewed articles published in English using the following search terms: (bird* OR avian*) AND (migrant* OR migratory) AND (agricultur* OR farm* OR agroforest*) AND (ecosystem service* OR crop pest* OR predation*). The search yielded a total of 287 hits. We screened all records to identify articles relevant to ecosystem services provided by migratory birds in agricultural systems within their wintering ranges. We then searched the reference lists of the initial set of articles for additional studies and screened the resulting publications, identifying six further articles and bringing the total number of reviewed articles to 293. In total, we identified 19 relevant articles (for details see Data S1).

## Appendix B. Sensitivity analyses accounting for differences in survey duration

**Table S1.**
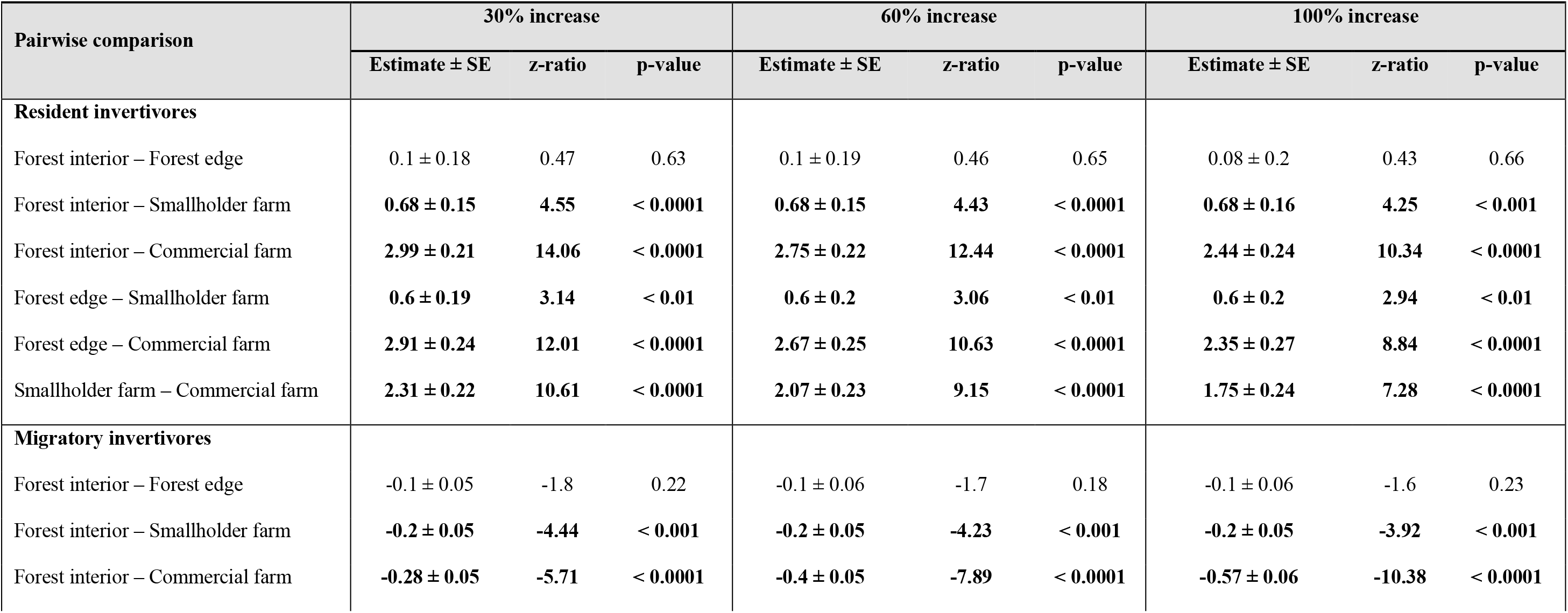
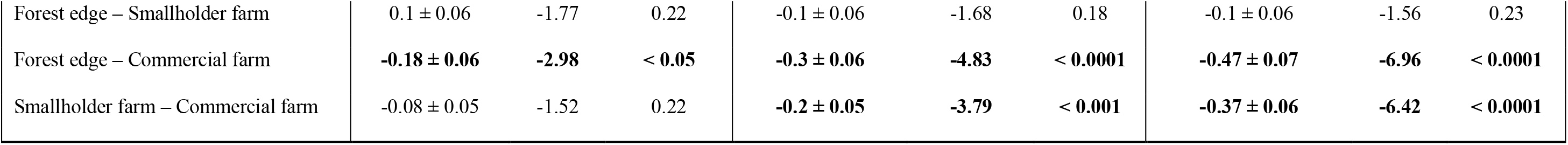
Sensitivity analyses of land-use effects on species richness under different survey-duration adjustments. Results show model summaries estimating the effects of land use on resident and migratory invertivore species richness. To account for shorter survey durations in commercial farms, species richness values for commercial farms were increased by 30%, 60% and 100% before modelling in three separate scenarios. Coefficients represent pairwise post hoc comparisons with Holm correction. Statistically significant effects are indicated in bold.

| **Pairwise comparison** | **30% increase** |  |  | **60% increase** |  |  | **100% increase** |  |  |
|---|---|---|---|---|---|---|---|---|---|
|  | **Estimate ± SE** | **z-ratio** | **p-value** | **Estimate ± SE** | **z-ratio** | **p-value** | **Estimate ± SE** | **z-ratio** | **p-value** |
| **Resident invertivores** |  |  |  |  |  |  |  |  |  |
| Forest interior – Forest edge | 0.1 ± 0.18 | 0.47 | 0.63 | 0.1 ± 0.19 | 0.46 | 0.65 | 0.08 ± 0.2 | 0.43 | 0.66 |
| Forest interior – Smallholder farm | **0.68 ± 0.15** | **4.55** | **< 0.0001** | **0.68 ± 0.15** | **4.43** | **< 0.0001** | **0.68 ± 0.16** | **4.25** | **< 0.001** |
| Forest interior – Commercial farm | **2.99 ± 0.21** | **14.06** | **< 0.0001** | **2.75 ± 0.22** | **12.44** | **< 0.0001** | **2.44 ± 0.24** | **10.34** | **< 0.0001** |
| Forest edge – Smallholder farm | **0.6 ± 0.19** | **3.14** | **< 0.01** | **0.6 ± 0.2** | **3.06** | **< 0.01** | **0.6 ± 0.2** | **2.94** | **< 0.01** |
| Forest edge – Commercial farm | **2.91 ± 0.24** | **12.01** | **< 0.0001** | **2.67 ± 0.25** | **10.63** | **< 0.0001** | **2.35 ± 0.27** | **8.84** | **< 0.0001** |
| Smallholder farm – Commercial farm | **2.31 ± 0.22** | **10.61** | **< 0.0001** | **2.07 ± 0.23** | **9.15** | **< 0.0001** | **1.75 ± 0.24** | **7.28** | **< 0.0001** |
| **Migratory invertivores** |  |  |  |  |  |  |  |  |  |
| Forest interior – Forest edge | -0.1 ± 0.05 | -1.8 | 0.22 | -0.1 ± 0.06 | -1.7 | 0.18 | -0.1 ± 0.06 | -1.6 | 0.23 |
| Forest interior – Smallholder farm | **-0.2 ± 0.05** | **-4.44** | **< 0.001** | **-0.2 ± 0.05** | **-4.23** | **< 0.001** | **-0.2 ± 0.05** | **-3.92** | **< 0.001** |
| Forest interior – Commercial farm | **-0.28 ± 0.05** | **-5.71** | **< 0.0001** | **-0.4 ± 0.05** | **-7.89** | **< 0.0001** | **-0.57 ± 0.06** | **-10.38** | **< 0.0001** |
| Forest edge – Smallholder farm | 0.1 ± 0.06 | -1.77 | 0.22 | -0.1 ± 0.06 | -1.68 | 0.18 | -0.1 ± 0.06 | -1.56 | 0.23 |
| Forest edge – Commercial farm | **-0.18 ± 0.06** | **-2.98** | **< 0.05** | **-0.3 ± 0.06** | **-4.83** | **< 0.0001** | **-0.47 ± 0.07** | **-6.96** | **< 0.0001** |
| Smallholder farm – Commercial farm | -0.08 ± 0.05 | -1.52 | 0.22 | **-0.2 ± 0.05** | **-3.79** | **< 0.001** | **-0.37 ± 0.06** | **-6.42** | **< 0.0001** |

**Table S2.** Sensitivity analyses of land-use effects on abundance under different survey-duration adjustments. Results show model summaries of land-use effects on resident and migratory invertivore abundance. To account for shorter surveys in commercial farms, species richness values there were increased by 30%, 60% and 100% before modelling. Coefficients represent pairwise post hoc comparisons with Holm correction.

| **Pairwise comparison** | **30% increase** |  |  | **60% increase** |  |  | **100% increase** |  |  |
|---|---|---|---|---|---|---|---|---|---|
|  | **Estimate ± SE** | **z-ratio** | **p-value** | **Estimate ± SE** | **z-ratio** | **p-value** | **Estimate ± SE** | **z-ratio** | **p-value** |
| **Resident invertivores** |  |  |  |  |  |  |  |  |  |
| Forest interior – Forest edge | -0.01 ± 0.1 | -0.01 | 0.99 | -0.01 ± 0.1 | -0.01 | 0.99 | -0.01 ± 0.1 | -0.01 | 0.99 |
| Forest interior – Smallholder farm | 0.19 ± 0.08 | 2.3 | 0.06 | 0.19 ± 0.09 | 2.23 | 0.08 | 0.19 ± 0.09 | 2.13 | 0.1 |
| Forest interior – Commercial farm | **1.15 ± 0.19** | **5.99** | **< 0.0001** | **0.98 ± 0.22** | **4.58** | **< 0.0001** | **0.77 ± 0.23** | **3.36** | **< 0.01** |
| Forest edge – Smallholder farm | 0.2 ± 0.11 | 1.83 | 0.14 | 0.19 ± 0.11 | 1.76 | 0.16 | 0.2 ± 0.11 | 1.68 | 0.19 |
| Forest edge – Commercial farm | **1.15 ± 0.2** | **5.68** | **< 0.0001** | **0.98 ± 0.23** | **4.37** | **< 0.001** | **0.78 ± 0.24** | **3.22** | **< 0.01** |
| Smallholder farm – Commercial farm | **0.96 ± 0.2** | **4.86** | **< 0.0001** | **0.79 ± 0.22** | **3.6** | **< 0.01** | 0.59 ± 0.24 | 2.49 | 0.05 |
| **Migratory invertivores** |  |  |  |  |  |  |  |  |  |
| Forest interior – Forest edge | -0.21 ± 0.28 | -0.76 | 0.45 | -0.21 ± 0.28 | -0.75 | 0.45 | -0.2 ± 0.28 | -0.73 | 0.47 |
| Forest interior – Smallholder farm | **-0.75 ± 0.21** | **-3.55** | **< 0.01** | **-0.73 ± 0.21** | **-3.47** | **< 0.01** | **-0.72 ± 0.21** | **-3.37** | **< 0.01** |
| Forest interior – Commercial farm | **-1.48 ± 0.28** | **-5.31** | **< 0.0001** | **-1.61 ± 0.3** | **-5.36** | **< 0.0001** | **-1.72 ± 0.3** | **-5.72** | **< 0.0001** |
| Forest edge – Smallholder farm | -0.54 ± 0.27 | -1.97 | 0.1 | -0.53 ± 0.27 | -1.93 | 0.11 | -0.52 ± 0.28 | -1.87 | 0.11 |
| Forest edge – Commercial farm | **-1.27 ± 0.33** | **-3.87** | **< 0.001** | **-1.41 ± 0.35** | **-4.05** | **< 0.001** | **-1.52 ± 0.35** | **-4.37** | **< 0.001** |
| Smallholder farm – Commercial farm | **-0.73 ± 0.27** | **-2.67** | **< 0.05** | **-0.88 ± 0.3** | **-2.96** | **< 0.01** | **-1.01 ± 0.3** | **-3.38** | **< 0.01** |

## Appendix C. Conceptual diagram illustrating the additional morphospace contributed by migrant invertivores.

**Figure S1.**
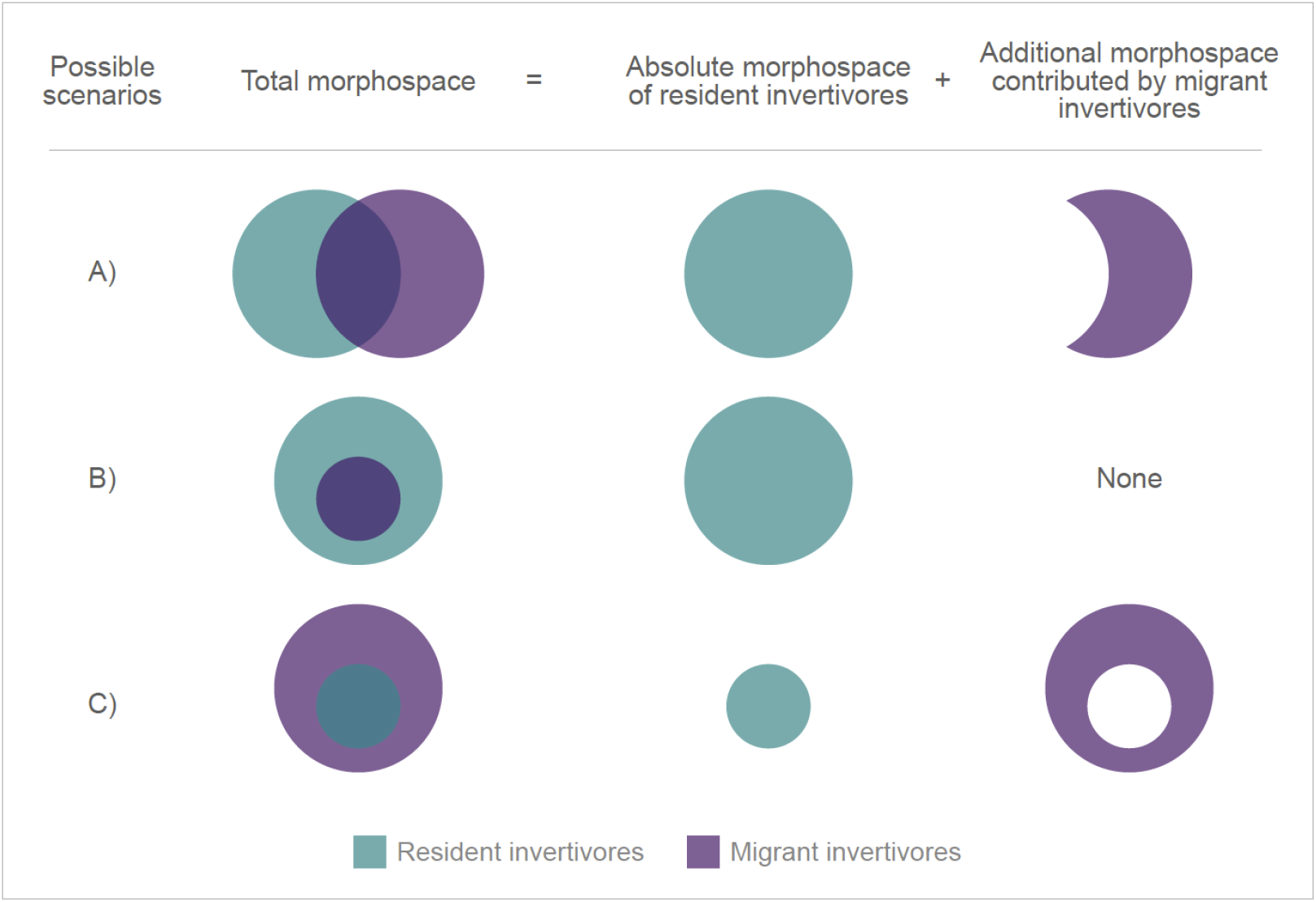
Conceptual diagram illustrating the partitioning of morphospace between resident and migratory invertivores and the additional morphospace contributed by migrants. Three possible scenarios are shown: A) Resident and migratory invertivore morphospaces partially overlap, with migrants contributing additional morphospace beyond that occupied by residents. B) Migratory invertivore morphospace lies entirely within resident invertivore morphospace, so migrants contribute no unique morphospace. C) Resident invertivore morphospace lies entirely within migratory invertivore morphospace, so migrants contribute the morphospace not occupied by residents. For ease of visualisation, the scenarios are illustrated in two dimensions. However, morphospaces were calculated using three beak morphometric measurements and should therefore be understood as three-dimensional, as shown in Figure 5.

## Appendix D. Results from pairwise post hoc tests

**Table S3.** Impact of land use on the species richness and abundance of resident and migratory invertivores. Results shown are model summaries of the pairwise post hoc tests investigating land-use effects on species richness and abundance of bird response groups. Statistically significant effects are indicated in bold. Effect sizes have not been back-transformed. For details on model structure, see Methods.

| **Pairwise comparison** | **Species richness** |  |  | **Abundance** |  |  |
|---|---|---|---|---|---|---|
|  | **Estimate ± SE** | **z-ratio** | **p-value** | **Estimate ± SE** | **z-ratio** | **p-value** |
| **Resident invertivores** |  |  |  |  |  |  |
| Forest interior – Forest edge | 0.09 ± 0.18 | 0.48 | 0.63 | -0.01 ± 0.01 | -0.01 | 0.99 |
| Forest interior – Smallholder farm | **0.68 ± 0.15** | **4.65** | **< 0.0001** | 0.2 ± 0.08 | 2.36 | 0.05 |
| Forest interior – Commercial farm | **3.23 ± 0.21** | **15.68** | **< 0.0001** | **1.39 ± 0.12** | **12.01** | **< 0.0001** |
| Forest edge – Smallholder farm | **0.6 ± 0.19** | **3.21** | **< 0.01** | 0.2 ± 0.11 | 1.88 | 0.12 |
| Forest edge – Commercial farm | **3.15 ± 0.24** | **13.39** | **< 0.0001** | **1.39 ± 0.13** | **10.5** | **< 0.0001** |
| Smallholder farm – Commercial farm | **2.55 ± 0.21** | **12.08** | **< 0.0001** | **1.2 ± 0.12** | **9.7** | **< 0.0001** |
| **Migratory invertivores** |  |  |  |  |  |  |
| Forest interior – Forest edge | -0.1 ± 0.05 | -1.88 | 0.25 | -0.21 ± 0.27 | -0.78 | 0.43 |
| Forest interior – Smallholder farm | **-0.2 ± 0.04** | **-4.65** | **< 0.0001** | **-0.77 ± 0.21** | **-3.67** | **< 0.01** |
| Forest interior – Commercial farm | **-0.15 ± 0.05** | **-3.26** | **< 0.01** | **-1.33 ± 0.27** | **-4.85** | **< 0.0001** |
| Forest edge – Smallholder farm | -0.10 ± 0.05 | -1.85 | 0.25 | -0.55 ± 0.27 | -2.04 | 0.11 |
| Forest edge – Commercial farm | -0.05 ± 0.06 | -0.92 | 0.65 | **-1.11 ± 0.32** | **-3.44** | **< 0.01** |
| Smallholder farm – Commercial farm | 0.05 ± 0.05 | 0.99 | 0.65 | -0.56 ± 0.29 | -2.09 | 0.11 |

**Table S4.**
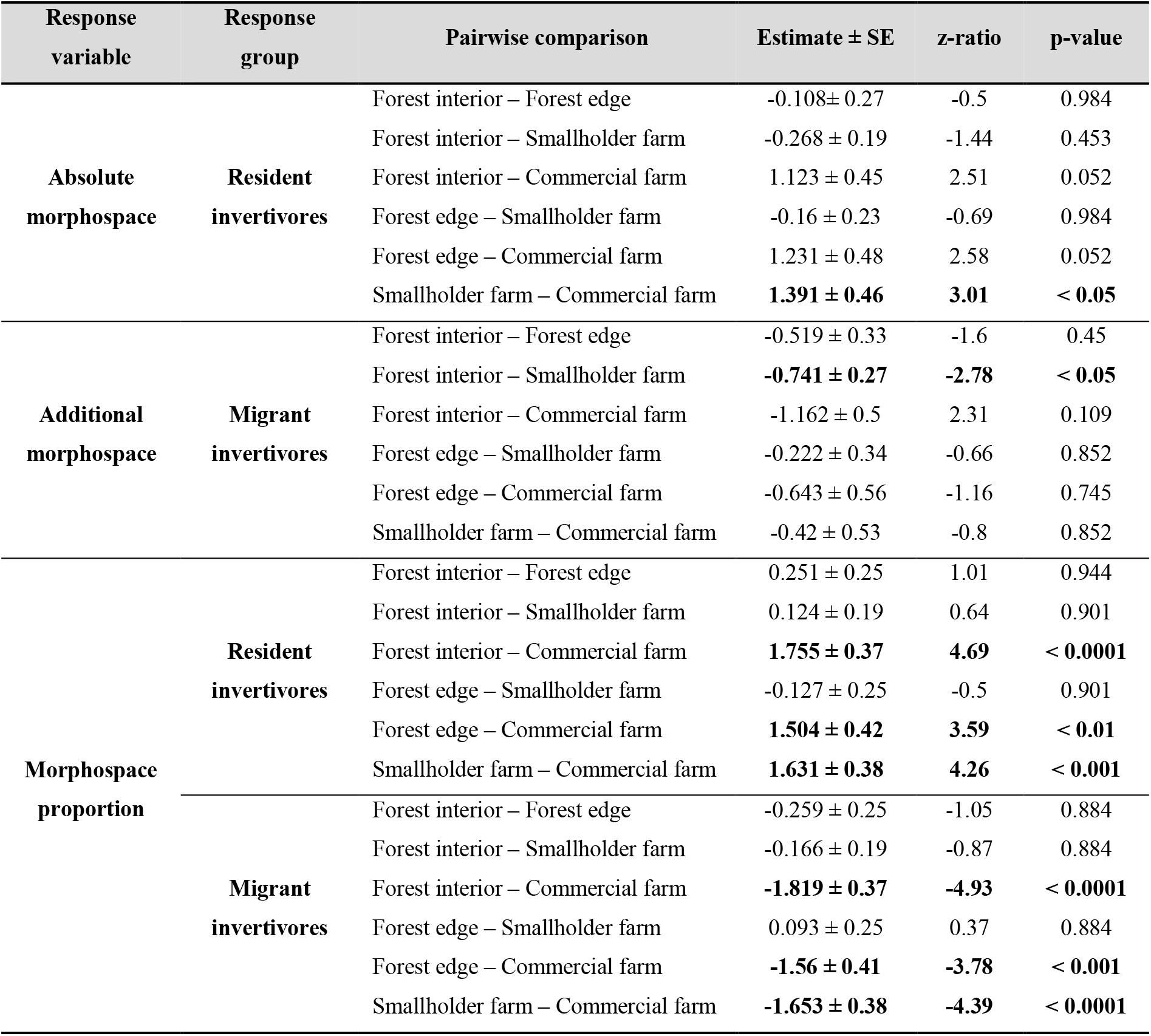
Impact of land use on the morphospace volume contributed by resident and migratory invertivores. Results shown are model summaries of pairwise post hoc tests investigating land-use effects on morphospace volume occupied by residents and the additional volume contributed by migrants (Figure S1). Coefficients are reported for absolute resident volume, additional migrant volume and the proportion of total volume occupied by each group. Statistically significant effects are indicated in bold. Effect sizes have not been back-transformed. For details on model structure, see Methods.

| Response<br>variable | Response<br>group | Pairwise comparison | Estimate ± SE | z-ratio | p-value |
|---|---|---|---|---|---|
| **Absolute<br>morphospace** | **Resident<br>invertivores** | Forest interior – Forest edge | -0.108± 0.27 | -0.5 | 0.984 |
|  |  | Forest interior – Smallholder farm | -0.268 ± 0.19 | -1.44 | 0.453 |
|  |  | Forest interior – Commercial farm | 1.123 ± 0.45 | 2.51 | 0.052 |
|  |  | Forest edge – Smallholder farm | -0.16 ± 0.23 | -0.69 | 0.984 |
|  |  | Forest edge – Commercial farm | 1.231 ± 0.48 | 2.58 | 0.052 |
|  |  | Smallholder farm – Commercial farm | **1.391 ± 0.46** | **3.01** | **< 0.05** |
| **Additional<br>morphospace** | **Migrant<br>invertivores** | Forest interior – Forest edge | -0.519 ± 0.33 | -1.6 | 0.45 |
|  |  | Forest interior – Smallholder farm | **-0.741 ± 0.27** | **-2.78** | **< 0.05** |
|  |  | Forest interior – Commercial farm | -1.162 ± 0.5 | 2.31 | 0.109 |
|  |  | Forest edge – Smallholder farm | -0.222 ± 0.34 | -0.66 | 0.852 |
|  |  | Forest edge – Commercial farm | -0.643 ± 0.56 | -1.16 | 0.745 |
|  |  | Smallholder farm – Commercial farm | -0.42 ± 0.53 | -0.8 | 0.852 |
| **Morphospace<br>proportion** | **Resident<br>invertivores** | Forest interior – Forest edge | 0.251 ± 0.25 | 1.01 | 0.944 |
|  |  | Forest interior – Smallholder farm | 0.124 ± 0.19 | 0.64 | 0.901 |
|  |  | Forest interior – Commercial farm | **1.755 ± 0.37** | **4.69** | **< 0.0001** |
|  |  | Forest edge – Smallholder farm | -0.127 ± 0.25 | -0.5 | 0.901 |
|  |  | Forest edge – Commercial farm | **1.504 ± 0.42** | **3.59** | **< 0.01** |
|  |  | Smallholder farm – Commercial farm | **1.631 ± 0.38** | **4.26** | **< 0.001** |
|  | **Migrant<br>invertivores** | Forest interior – Forest edge | -0.259 ± 0.25 | -1.05 | 0.884 |
|  |  | Forest interior – Smallholder farm | -0.166 ± 0.19 | -0.87 | 0.884 |
|  |  | Forest interior – Commercial farm | **-1.819 ± 0.37** | **-4.93** | **< 0.0001** |
|  |  | Forest edge – Smallholder farm | 0.093 ± 0.25 | 0.37 | 0.884 |
|  |  | Forest edge – Commercial farm | **-1.56 ± 0.41** | **-3.78** | **< 0.001** |
|  |  | Smallholder farm – Commercial farm | **-1.653 ± 0.38** | **-4.39** | **< 0.0001** |

